# Actigraphic Correlates of Neuropsychiatric Disability in Adult Patients with Focal Epilepsy

**DOI:** 10.1101/2022.10.18.512750

**Authors:** Mark A. Abboud, Jessica L Kamen, John S Bass, Lu Lin, Jay R. Gavvala, Sindhu Rao, Stephen F Smagula, Vaishnav Krishnan

**Affiliations:** Departments of Neurology, Baylor College of Medicine, Houston, TX USA; Departments of Psychiatry and Epidemiology, University of Pittsburgh Medical Center, Pittsburg PA USA; Neuroscience, Baylor College of Medicine, Houston, TX USA; Psychiatry & Behavioral Sciences, Baylor College of Medicine, Houston, TX USA

## Abstract

Disability in patients with epilepsy (PWE) is multifactorial: beyond seizure frequency/severity, PWE are prone to a range of neuropsychiatric, cognitive, and somatic comorbidities that significantly impact quality of life. In this study, we explored how variations in epilepsy severity and the burden of self-reported somatic/neuropsychiatric symptoms are associated with disruptions to 24h activity patterns (rest-activity rhythms, RARs), determined through wrist accelerometry/actigraphy. Continuous multiday recordings were obtained from 59 adult patients with focal epilepsy (44% male, ages 18-72), who contemporaneously provided responses to a range of validated psychometric instruments to measure the burden of anxiety, depression, sleepiness, and somatic symptoms. As a comparator, we conducted a similar psychometric-actigraphic correlation in 1761 subjects of Hispanic origin (35% male, ages 18-65) from the Study of Latinos (SOL) Sueño Ancillary Study. RARs were analyzed via a sigmoidally-transformed cosine model (quantifying RAR amplitude, steepness, acrophase and robustness) and non-parametric measures to estimate RAR stability, fragmentation, and sleep. Compared with age- and sex-matched SOL subjects, RARs from PWE subjects featured a significantly diminished amplitude, a wider rest phase and significantly more total daily sleep. Within PWE, similar RAR distortions were associated with seizure intractability and/or anticonvulsant polytherapy. In contrast, high anxiety, depression, and somatic symptom scores were associated with diminished RAR robustness and a delayed acrophase. We applied the complete SOL Sueño database to train logistic regression models to dichotomously classify anxiety, depression and sleepiness symptoms using age, sex, body mass index and a range of non-collinear RAR parameters. When tested on PWE, these models predicted prevalent anxiety and depression symptoms with modest success (accuracy ∼70%) but failed to predict subjective sleepiness. Together, these results demonstrate that RAR features may vary with depression and anxiety symptoms in ambulatory patients with focal epilepsy, potentially offering a set of objective wearable-derived endpoints to adjunct routine clinical care and drug/device treatment trials. With larger actigraphic-psychometric datasets in PWE, we may identify RAR signatures that can more precisely distinguish between variations in seizure risk, the burden of anticonvulsant therapy and prevalent mood/anxiety symptoms.

## Introduction

Across a range of chronic medical conditions, an exciting and growing body of research seeks to define how wearable devices may be beneficial to monitor symptom severity, treatment compliance and/or response. Some of these technologies have already substantially impacted patient care (e.g., smart watch EKG monitors to detect atrial fibrillation, implanted glucose sensors for diabetic patients). The bulk of wearable science in epilepsy research has focused on the problem of seizure detection/forecasting by applying machine learning to annotated recordings of continuous physiological (e.g., electrodermal activity, heart rate, temperature) and/or neurophysiological biomarkers (e.g., ambulatory electrocorticography)^1^. Relatively fewer resources have been devoted to understanding whether wearables may inform the management of the many psychiatric and somatic comorbidities of epilepsy^2^. Adults with PWE are frequently impacted by neuropsychiatric symptom constellations that include features of depression, anxiety, insomnia/excessive sleepiness and subjective/objective memory impairment^3-8^. Several lines of evidence indicate that the severity of comorbid psychiatric symptoms generally correlate with markers of seizure severity/intractability^9-13^. Nevertheless, many patients experience persistent neuropsychiatric symptoms despite sustained seizure control, highlighting the complex and multifactorial determinants of mental wellbeing in PWE. Antiseizure medications (ASMs) are one such complex determinant: psychiatric and behavioral side effects vary significantly among ASMs, but are also more frequently experienced by patients with medically refractory/intractable epilepsy and/or pre-existing psychiatric disease^14,15^, the latter itself a risk factor for seizure intractability^16,17^.

Regardless of etiology, early detection and interventions designed to address these frequently undetected and undertreated comorbidities can provide tangible health-care benefits for the patient and their caregivers^2,18^. To this end, several psychometric screening tools/instruments have been applied and validated in PWE, including the PHQ-9 (Patient Health Questionnaire-9 [depression])^19^, the GAD-7 (Generalized Anxiety Disorder-7)^20^ and the ESS (Epworth Sleepiness Scale)^21^. These instruments generally lend fluidly to routine patient care and formal research endeavors, given their brevity (7-9 questions), availability in multiple languages^22-24^ and quantitative endpoints that conveniently facilitate linear or logistic regression analyses. Their limitations must also be recognized. First (pertaining to *some* PWE), they assume intact language function, and are therefore nearly impossible to reliably administer in patients who are aphasic or nonverbal. Whether more subtle verbal perception deficits (associated with dominant temporal lobe epilepsies/temporal lobectomies^25^) may impact self-report remains poorly explored. Second, unlike measurements of body weight or glycated hemoglobin, psychometric surveys are prone to conscious or unconscious response distortion that may result in over- or under-reporting of psychiatric symptoms^26,27^. Under-reporting may be particularly common in patients that are already experiencing the societal stigma associated with an epilepsy diagnosis. Further, while many questionnaires remind subjects to focus on a coarse timescale (“During the last 2 weeks, how often have you …”), responses may be distorted by more acute stressors, including recent seizures. Third, despite claims of test-retest reliability, these instruments were not designed for frequent repeated measurements, but rather as an initial screening tool to document symptom severity and consider psychiatric consultation/initiate psychopharmacological treatments. This raises questions about whether “trends” in survey responses collected over months/years accurately reflect variations in symptom burden. And finally, many different forms of these surveys are available and are inconsistently used across studies. As one example, a recent systematic electronic database review identified at least *20* different psychometric instruments that have been validated to assess anxiety in studies emerging from low to middle income countries^28^.

Digital phenotyping tools provide objective, continuous and longitudinal solutions to monitor symptom severity that may complement the subjective, clinic-based appraisals of neuropsychiatric symptomatology^29^. Among the many digital endpoints that have been explored (including facial expression, speech analysis, pose estimation), efforts employing wrist-worn accelerometers (“wrist-actigraphy”) have been most rigorously investigated. While initially conceived to monitor the quantity and timing of sleep, continuous several day-long actigraphy data can capture changes in 24h patterns of activity, termed rest-activity rhythms (RARs). RAR distortions have been described and meta-analyzed for studies employing subjects with major depression^30^, bipolar^31^ and anxiety disorders^32^. Among parametric techniques designed to analyze/model RARs, standard cosine approaches have now been largely replaced by sigmoidally transformed cosine models^33^ (“extended cosinor analysis/ECA”), to accommodate the more *square* features of human RARs. ECA provides quantitative measures of RAR morphology (amplitude, steepness, rest/activity ratios and the timing of peak activity [acrophase]) and RAR robustness (F, or *pseudo F-statistic*), where lower F values indicate a more erratic or irregular RAR. The ECA technique has been most widely applied in cross-sectional case-control studies of depression: multiple published case-control studies have shown that prevalent depression symptoms are associated with a lower F^34-37^ and amplitude^34,36^. In a recent analysis of 1800 older individuals (mean age 72.9 years) from the NHANES (National Health and Nutrition Examination and Survey) accelerometer study, prevalent depression symptoms (PHQ-9 ≥ 10) *and* mild cognitive impairment were enriched in subgroups that displayed a delayed acrophase and low F^38^. RARs also vary significantly with age, sex and body habitus. A recent published analysis of the larger NHANES accelerometer dataset (8200 subjects ≥20 years of age) found that females display *higher* RAR amplitudes and F values across the age spectrum, while also reporting clear age-dependent decreases in amplitude, increases in F and a left-shifted (earlier) acrophase^39^. Obesity, as defined by the body mass index (BMI), was also shown to be associated with *lower* amplitudes and F^40^.

In this study, our objective was to describe how variations in seizure severity, ASM burden and neuropsychiatric comorbidity may associate with RAR distortions in an initial cohort of 59 adult patients with focal epilepsy. In addition to continuous actigraphic recordings, subjects were asked to complete a set of widely utilized psychometric instruments for depression (PHQ9, QIDS-SR [Quick Inventory of Depression Symptoms – Self Report]), anxiety (GAD-7), sleepiness (ESS) and somatic symptoms (AEP [Adverse Event Profile^41,42^, PHQ-15^43^). We utilized the ECA technique to quantify RAR morphology and applied a simple algorithm to estimate the timing and duration of sleep, allowing us to understand whether self-reported measures of subjective sleepiness (ESS) at all correlate with patterns of sleep timing and occurrence. In place of a contemporaneously measured “healthy control” population, we conducted a similar actigraphic-psychometric correlation from ∼1700 subjects of Hispanic origin that participated in the Study of Latinos (SOL) ^44,45^. These individuals wore a similar actigraphy device, over a similar number of days, and similarly underwent psychometric assessments of subjective anxiety, depression and sleepiness. From this SOL dataset, we derived models to predict symptom prevalence dichotomously (absent vs affected), based on age, sex and RAR characteristics, and tested the accuracy of this model in subjects with epilepsy.

## Methods

### Participants and Study Design

All study protocols were approved by the Baylor College of Medicine Institutional Review Board. Subjects were recruited and consented from the Baylor Comprehensive Epilepsy Clinic (Houston, TX) during routine follow up visits. We included English-speaking patients, aged 18-75, with partial onset seizures as determined by reported/recorded ictal semiology and/or documented interictal or ictal EEG data. We excluded (i) pregnant patients, (ii) those with cognitive impairment/intellectual disability severe enough to preclude self-report or self-consent, and (iii) those with significant mobility limitations (e.g., hemiparesis, quadriparesis). Subjects were instructed to wear the rugged, widely popular^46-51^ and FDA-approved Actiwatch-2 device (Phillips Respironics) continuously for at least 10 days on their nondominant wrist (“through sleep and showers”), and asked to complete a printed handout of psychometric surveys (QIDS-SR, ESS, AEP, PHQ-<u>SAD</u>S [which includes the PHQ15-somatic, GAD7-<u>a</u>nxiety and PHQ9-<u>d</u>epression]). A prepaid envelope was provided to return study materials.

### Study of Latinos (SOL)

The Hispanic Community Health Study/Study of Latinos (HCHS/SOL) enrolled *16415* non-institutionalized self-identified Hispanic/Latino adults (aged 18-74), from 9872 randomly selected households in Miami, Bronx, San Diego and Chicago^52^. The main goals of this prospective cohort study were to characterize risk and protective factors for a range of chronic conditions. An initial comprehensive baseline assessment for all subjects (2008-2011) was followed by an ancillary actigraphy study (“SUEÑO”, 2010-2013) for 2252 subjects belonging to the greater SOL cohort, aged 18-65. Sueño exclusion criteria included current pregnancy, a diagnosis of narcolepsy, sleep apnea or previous/current CPAP/BIPAP treatment. Since subjects were not specifically asked about a current or historical diagnosis of epilepsy/seizures, we assume that the prevalence of epilepsy within these subjects likely mirrors the general population (∼0.8-1%)^53^. Participants were instructed to wear an Actiwatch Spectrum Device (Phillips Respironics) for a week and were administered a range of wellbeing surveys over the phone in their choice of English or Spanish. Our analysis focused on the following measures: ESS, STAI-10 (10 item State Trait Anxiety Scale^54^), and the CESD-10 (10-item Center for Epidemiologic Studies Depression Scale^55^). Subjects were also administered the HSI (Hispanic Stress Inventory^56^) and the NSC (Neighborhood Social Cohesion Scale^57^). 1887 participants consented to allow their data for public use: their demographic data, psychometric scores and raw actigraphy data were downloaded from the National Sleep Research Resource (www.sleepdata.org). We further excluded 24 subjects for whom BMI or ESS/CESD-10/STAI-10 data were unavailable, 2 subjects that displayed an excessive number of missing actigraphy data (>200 minutes/day), and 100 patients in whom an extended cosinor analysis returned a non-physiological solution (e.g., amplitude < mesor [77], impossible fit [15] or poor circadian rhythmicity resulting in an alpha = ± 1 [9]). Age, gender, BMI and ECA trends for the remaining 1761 subjects are summarized in Supplemental Figure 1. The median number of complete days of recording was 6 (range 4-16). For the case-control comparison shown in Fig. 1D-F, every epilepsy patient (except for two patients aged > 65) was matched to three age- and gender-matched SOL subjects. Due to the chance distribution of available subjects in the SOL dataset, three patients with epilepsy (1598527, 6784058, 8600154) were paired with gender-matched SOL subjects who were a year older or younger. For many age/sex combinations, there were many more than three matches were available (e.g., SOL contains at least nine 23-year old female subjects), we chose subjects with the three most median F values.

**Figure 1:**
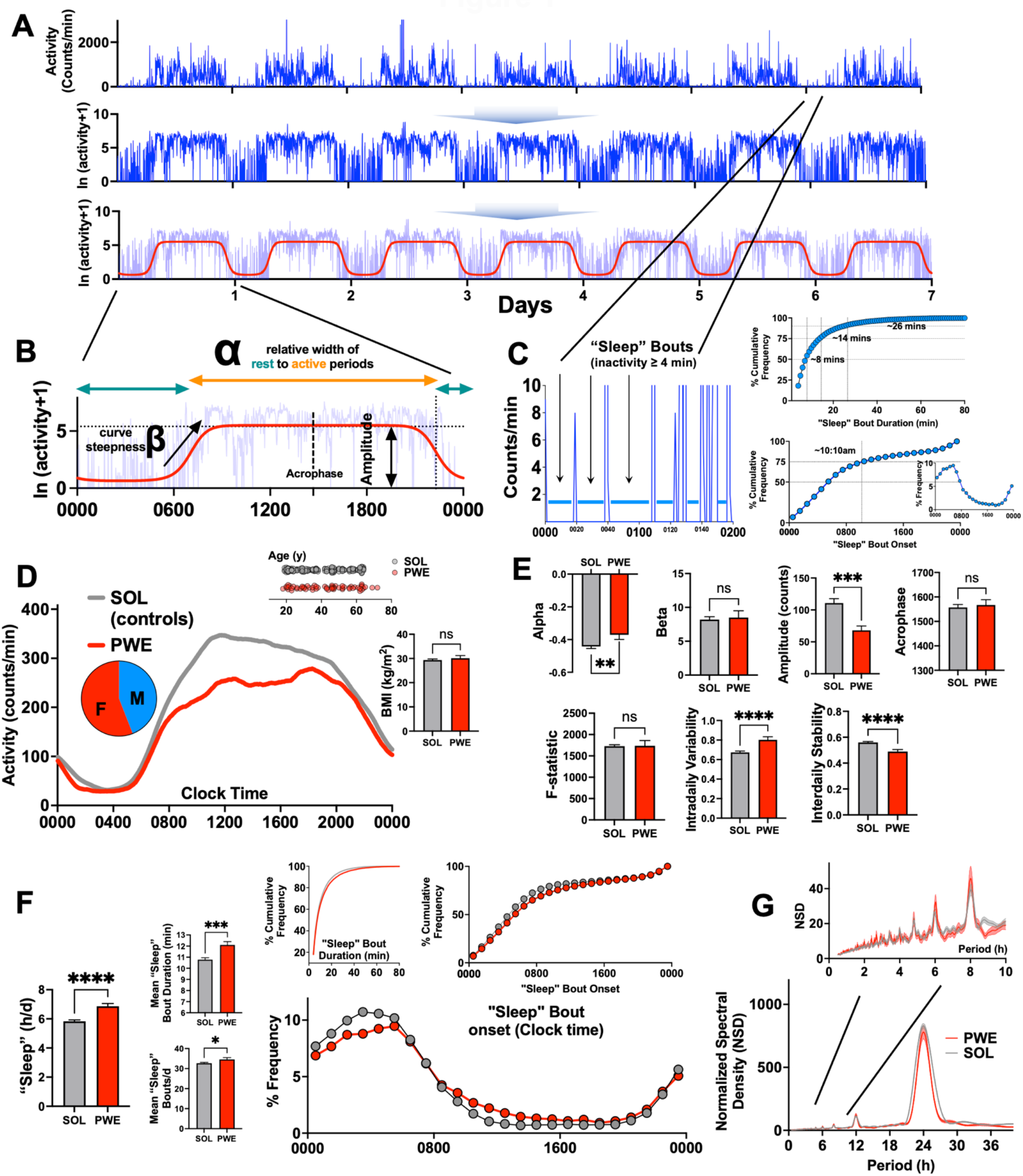
RAR Characteristics in Patients with Focal Epilepsy. A. Example of a 7-day patient actogram, log-transformed and least squares regressed to a sigmoidally extended cosine function. B. Parameters extracted include alpha, beta, amplitude, acrophase and F (or *pseudo-F* statistic, not shown). C. Approach to identifying bouts of “sleep”, with standard and cumulative distributions of “sleep” bouts by duration and start times. D. Averaged actogram from PWE and age- and sex-matched SOL controls subjects, with similar average body mass index. E. Compared with SOL controls, PWE displayed a significantly lower amplitude and higher alpha (indicating a wider rest phase). PWE also displayed a relative increase in intradaily variability and decrease in interdaily stability. F. PWE accumulate greater total daily “sleep”, through a combination of longer (KS test, p <0.0001) and more frequent “sleep” bouts. “Sleep” bouts in PWE are more widely dispersed throughout the day. G. Spectral decomposition of raw actigraphy signal demonstrating a similar major peak (circadian oscillator, ∼24h) and ultradian peaks (inset). Mean ± s.e.m shown for all. *, **, ***, **** indicating p < 0.05, p<0.01, p<0.001, p<0.0001 respectively (unpaired two-tailed Student’s T test).

### Actigraphy, Extended Cosinor Analysis, Sleep

The Actiwatch Spectrum and Actiwatch-2 devices utilize piezoelectric sensors to measure unidimensional accelerometry, with the sensor oriented to detect natural movements of the wrist. At 32Hz, an analog voltage signal is digitized to obtain a running baseline value and constant gravity-related acceleration is filtered out. For each second, the maximum deviation from the running baseline is logged and summed into either 15, 30 or 60s epochs (user-selected, with longer epochs prolonging battery life). SOL data were collected in 30s epochs, which were down-sampled to 60s epochs (by adding consecutive values) to compare with data from epilepsy subjects (60s epochs). Actograms from both cohorts were visually inspected for periods of non-watch wearing. Incomplete days were excluded. RAR parameters were quantified as previously described^33,38,39^ using an open-source R package (https://github.com/JessLGraves/RAR). We focused on the following parameters: alpha (relative width of rest to active phase), beta (measuring sigmoid steepness), acrophase (midpoint of the active period), amplitude (fitted curve maximum), F (or *pseudo F-statistic*, proportional to RAR robustness), IS (interdaily stability) and IV (intradaily variability)^58^. To estimate “sleep” (enquoted to emphasize its derivation from purely actigraphic measures), we identified the timing and length of bouts of zero activity lasting ≥ 4 minutes (Fig. 1C). This approach models a popular neurophysiologically validated algorithm employed to noninvasively estimate sleep in rodent species^59^. Unlike existing models that employ moving averages to ignore brief awakenings^60^, this approach recognizes such awakenings as true interruptions of sleep^61^. It does not require sleep diaries, makes no assumptions about a “main sleep period”^62^ and is agnostic to sleep stage. “Sleep” bouts identified in this manner (from PWE subjects) displayed a median duration of 8 minutes, with 75% of all such “sleep bouts” occurred between 0000 (midnight) and 1010hrs (Fig. 1C). In the Sueño dataset (n=1761), this approach identified a median daily “sleep” duration of ∼5.7h/d (standard deviation 1.2h), similar to but lower than previous estimates (using the same dataset) that incorporated diary data and event markers (6.7 ± 1.1h)^62,63^.

### Chronic Intracranial EEG Review

Four subjects were implanted with responsive neurostimulation devices (RNS)^64^ during actigraphic recordings, three of whom were receiving active close-loop stimulation. Neurostimulation parameters remained unchanged during the study period. Hourly measures of tailored neurostimulation bursts (treatments) and long episodes (sustained bursts of epileptiform activity that may or may not be associated with a clinical correlate) were tallied through the NeuroPace Patient Data Management System (PDMS), a secure online interface. Concurrent actigraphy and RNS data are plotted in Figure 5 for all three subjects.

### Statistical Analysis

Graphing and statistical analysis was conducted on Prism 9. Representative activity curves (e.g., Fig. 1D) were tallied by plotting the minutely arithmetic mean of actigraphy counts for all patients across all days. This mean curve was then smoothed by a 2^nd^ order polynomial averaging 100 neighboring points on either side. Unpaired two-tailed Student’s t tests were applied to compare two groups (Fig. 1D), one-way analysis of variance for > 2 groups (e.g., Fig. S1D) or two-way analysis of variance when two independent variables were examined (e.g., Fig. S1C). Raw distributions were compared by Kolmogorov-Smirnov (KS) testing (e.g., “sleep” bouts, Fig. 1F). Chi-squared analyses to compare occurrences were always two-tailed (without the Yates correction). Lomb-Scargle periodograms (MATLAB) were calculated to measure peaks and power of circadian/ultradian rhythms (Fig. 1G). Exploratory analyses are depicted in Fig.2 and 3, utilizing either dichotomous variables (e.g., sex, prior epilepsy surgery) or continuous variables separated by a median split (see Table 1 for medians). We defined polytherapy as the concurrent intake of ≥2 daily ASMs; “as needed”/rescue benzodiazepine prescriptions were not considered ASMs. Unless otherwise specified, mean ± standard error of the mean is depicted. Multiple logistic regression was performed on the SOL study set to individually model survey responses for ESS (median 5), STAI-10 (median 15) and CESD-10 (median 6) as binary variables (LOW = median or less, HIGH > median). Multicollinearity was estimated with variance inflation factors (VIF), and variables with a VIF > 4 were excluded. One categorical (sex) and 9 continuous Z-normalized predictor variables were employed (age, BMI, total “sleep”/day, alpha, beta, amplitude, F, acrophase, # days of recording). The same variables for epilepsy subjects (Z-normalized separately) were applied to measure model accuracy. K-means clustering (MATLAB) was performed with 2000 replicates to identify the solution with the lowest total sum of Euclidean distances.

**Figure 2:**
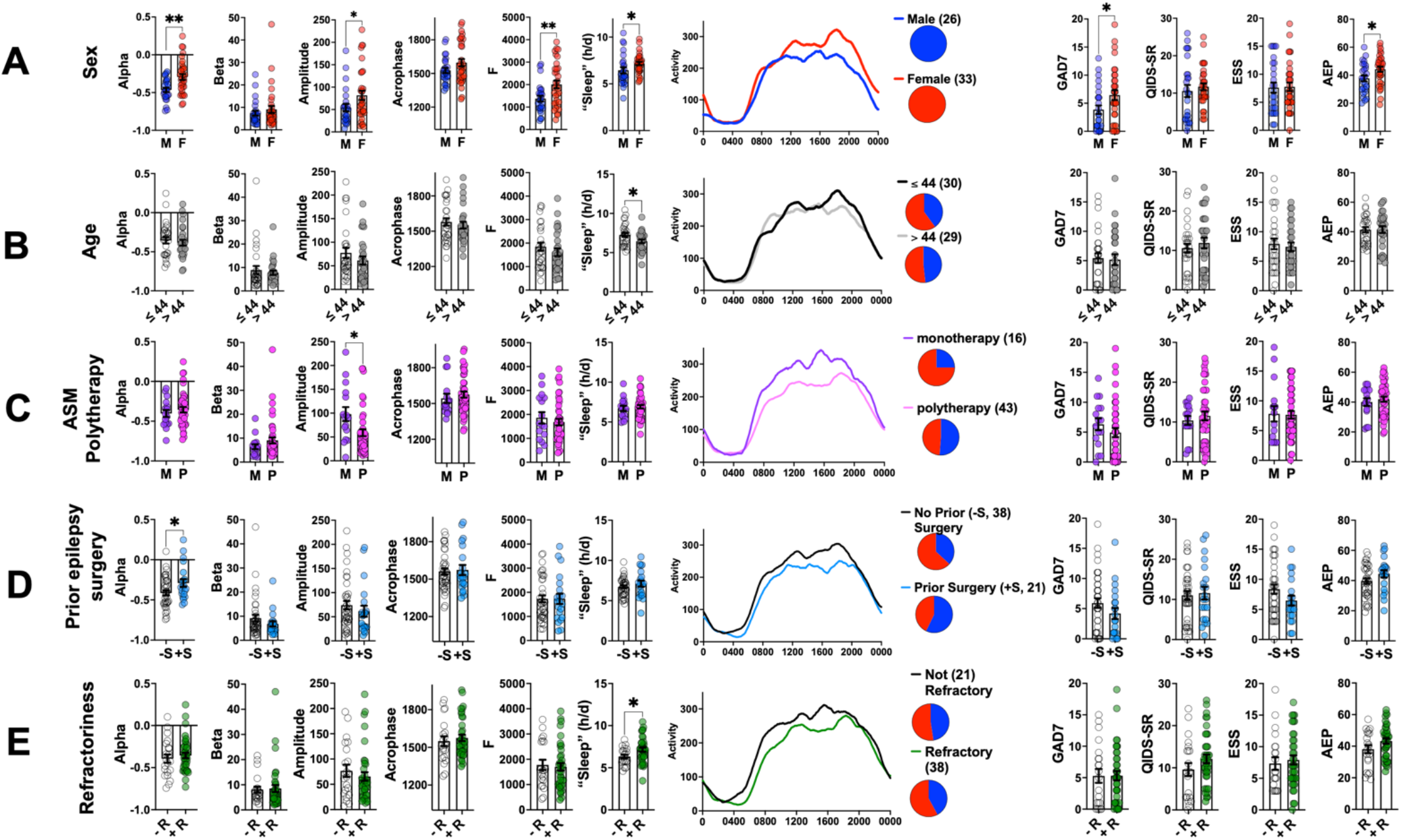
RAR Variations by Sex, Age and Markers of Epilepsy Severity. Pairwise comparisons of actigraphic (left) and psychometric survey responses (right), with average smoothed actograms (center). Mean ± s.e.m shown for all. *, ** indicated p < 0.05, p<0.01 respectively (unpaired two-tailed Student’s T test). ESS: Epworth Sleepiness Scale, AEP: Adverse Event Profile, PHQ: Patient health questionnaire, GAD-7: Generalized anxiety disorder-7, QIDS-SR: Quick Inventory of Depression-Self Report.

**Figure 3:**
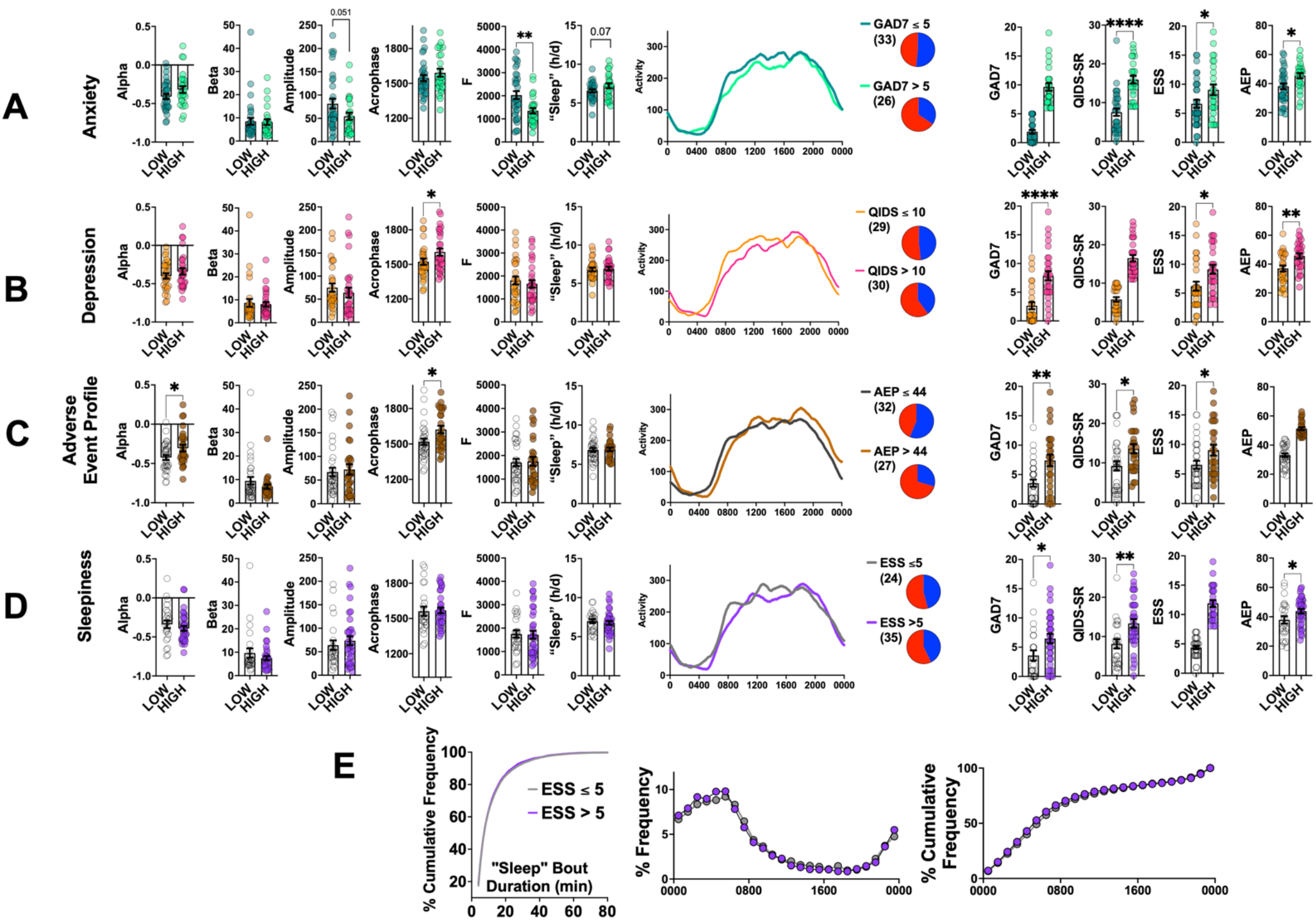
RAR Variations by Neuropsychiatric Symptomatology. Pairwise comparisons of actigraphic (left) and psychometric survey responses (right), with average smoothed actograms (center). Groups were separated by a median split, including scores on GAD-7 (A), QIDS-SR (B), AEP (C) and ESS (D), with median scores shown in Table 1. E: Frequency distributions for sleep bout durations (KS test, p>0.1) and timing comparing low ESS to high ESS groups. Mean ± s.e.m shown for all. *, **, **** indicating p < 0.05, p<0.01 and p<0.0001 respectively (unpaired two-tailed Student’s T test). ESS: Epworth Sleepiness Scale, AEP: Adverse Event Profile, PHQ: Patient Health questionnaire, GAD-7: Generalized anxiety disorder-7, QIDS-SR: Quick Inventory of Depression-Self Report.

**Table 1:** Demographic and clinical data of PWE subjects. BMI: Body mass index, ASM: antiseizure medication, MRI: magnetic resonance imaging, VNS: vagal nerve stimulation, RNS: responsive neurostimulation, SSRI: Selective serotonin reuptake inhibitors, LEV: levetiracetam, ESS: Epworth Sleepiness Scale, AEP: Adverse Event Profile, PHQ: Patient health questionnaire, GAD-7: Generalized anxiety disorder-7, QIDS-SR: Quick Inventory of Depression-Self Report.

| Variable | # (%) or median (range) |
| --- | --- |
| Sample Size | 59 |
| Female Sex | 33 (55.9%) |
| # Days of Recording | 7 (4-12) |
| Age | 44 (18-72) |
| BMI (kg/m<sup>2</sup>) | 28.3 (16.1-53.1) |
| Employed/ in school | 27 (44.8%) |
| Refractory Epilepsy | 38 (64.4%) |
| #ASMs | 2 (1-5) |
| On ASM Monotherapy | 16 (27%) |
| Temporal Lobe Onset<br>(or suspected temporal lobe onset) | 37 (62.7%) |
| Normal MRI | 30 (50.8%) |
| Active VNS Therapy | 6 (10.1%) |
| Prior ATL | 5 (8.4%) |
| Active RNS Therapy | 3 (5.1%) |
| Concurrent SSRI Use | 14 (23.7%) |
| Concurrent Antipsychotic Use | 3 (5.1%) |
| LEV intake | 27 (45.8%) |
| ESS | 7 (0-19) |
| AEP | 43 (19-63) |
| PHQ-15 | 7 (0-21) |
| PHQ-9 | 7 (0-27) |
| GAD-7 | 5 (0-19) |
| QIDS-SR | 11 (0-26) |

## Results

### RAR Characteristics in PWE and SOL

67 eligible subjects with focal epilepsy consented. Data from 8 subjects were not included: 3 patients never returned the study packet (of which 2 were permanently lost to follow up), 1 patient’s actigraphy device experienced a technical malfunction, 2 patients returned the packet without ever wearing the watch, and an additional 2 patients provided <4 full days of watch wearing. Demographic and epilepsy-related variables for the remaining 59 patients included in the study are summarized in Table 1 and elaborated in Supplemental Table 1. As might be expected from a tertiary epilepsy referral clinic, almost 2/3^rds^ of our sample displayed medically intractable epilepsy with >70% receiving ASM polytherapy. ∼1/3^rd^ of patients had received some form of epilepsy surgical treatment, including vagal nerve stimulation (∼10%), responsive neurostimulation (∼5%) and prior anterior temporal lobectomies (∼8%).

RAR parameters for all PWE subjects were compared against three age- and gender-matched SOL Sueño subjects, selected from a larger cohort of 1761 participants which were ∼65% female with a median age of 49 (range 19-65, Supplemental Figure 1A, B). In comparison to these SOL matched subjects, PWE displayed a significantly lower amplitude and narrower active phase (Fig. 1D, E). Overall measures of RAR robustness (F) were similar between groups, although PWE displayed greater within-day rhythm fragmentation (IV or intradaily variability) and diminished across-day similarity (IS or interdaily stability). Similar IV and IS changes were observed in one previous actigraphic evaluation of patients with epilepsy and healthy controls (n=17-22)^65^, which did not employ ECA or concurrent psychometric surveys.

The timing and duration of bouts of “sleep” were estimated by tallying epochs of sustained immobility lasting ≥ 4 minutes (see Methods). PWE accumulated greater total daily “sleep”, associated with more frequent and slightly more prolonged “sleep” bouts (Fig. 1F). Compared with SOL subjects, “sleep” bouts in PWE were more widely dispersed across the 24h cycle. To reveal any gross disturbances in rhythmicity, we spectrally decomposed raw actigraphy time series data across both cohorts (Fig. 1G). Both groups demonstrated a dominant peak at ∼24h (circadian oscillator), and a similar distribution of ultradian activity rhythms.

We then examined whether RAR parameters within the larger SOL cohort vary by age, gender and BMI (as previously reported within NHANES actigraphy datasets^39,40^). While we did not identify clear sex- or age-dependent changes in RAR amplitude, we identified a clear increase in F scores in female subjects and an age-dependent *left*-shifted acrophase (earlier mid-point, Fig. S1C). Female subjects also displayed a higher alpha level and accumulated significantly more total daily “sleep”, consistent with at least one previous cross-sectional actigraphic study of 669 subjects aged 38-50^66^. These results are also generally aligned with estimates from sleep diaries^67^, and further validate our algorithm. In a univariate analysis, stepwise increases in BMI were also associated with significantly lower F scores and RAR amplitudes (Fig. S1D), replicating findings from the NHANES cohort^40^. These SOL results, while uncontrolled for a range of health variables, re-emphasize the importance age, sex and BMI in assessing RAR morphology.

### RAR changes within PWE

We next explored how RAR changes and psychometric parameters varied within the PWE sample by examining a series of pair-wise comparisons. Our goal was to achieve a descriptive overview and explore whether variations in seizure severity and neuropsychiatric comorbidity may be linked to unique or overlapping RAR distortions. Without formally correcting for multiple comparisons, we recognize that at least 5% of significant differences may be spurious. With these caveats in mind, sex differences in RAR parameters within PWE nicely aligned with SOL (Fig. S1C) and NHANES results^39^, with female subjects displaying a significantly higher F, amplitude and alpha (∼wider rest phase). Female subjects also displayed a significant increase in GAD-7 and AEP scores (Fig. 2A). Older patients (>44 [median]) displayed significantly lower total daily “sleep” (Fig. 2B). To approximate RAR changes that vary by epilepsy severity, we explored the role of three distinct but related surrogate measures. ASM polytherapy was associated with a significant reduction in amplitude (Fig. 2C), while a history of prior epilepsy surgery (including neuromodulation devices) was associated with a higher alpha (wider rest phase, Fig. 2D). Subjects with medically intractable epilepsy (refractoriness) displayed significantly greater total daily “sleep” (Fig. 2E). Through either lens, subjective measures of anxiety, depression, daytime sleepiness, or somatic symptoms were not significantly different in patients with greater epilepsy severity. Across all five comparisons, BMIs were not significantly different. Similarly, aside from age-related changes (Fig. 2B), mean subgroup ages were also not significantly different (not shown).

In an analogous tactic, we examined how variations in psychometric survey scores were associated with changes in RAR measures. As a first pass, we divided patients into two groups (low vs high) through a median split. This approach revealed symptom multimorbidity, such that patients assigned to the high GAD-7 group (as an example) also displayed significantly greater scores on the QIDS-SR, ESS and AEP (Fig. 3). However, specific symptom clusters revealed varied RAR changes. High GAD-7 scores were associated with a significantly lower F (Fig. 3A), while high QIDS-SR scores were associated with a delay in acrophase (Fig. 3B). Subjects with high AEP scores displayed both a delay in acrophase and larger alpha (Fig. 3C). Finally, patients with greater than median ESS scores displayed RAR parameters that were not significantly different from low-ESS patients (Fig. 3D). Interestingly, these groups were largely indistinguishable in terms of total daily “sleep”, “sleep” bout durations or timing (Fig. 3E). Across all four comparisons, BMIs and ages were not significantly different (not shown) except for AEP (Fig. 3C), where subjects with high AEP scores had a greater mean BMI (33.1 vs 27.5, p<0.01).

### Can RAR parameters predict prevalent comorbid psychiatric symptoms?

In the final step, we explored actigraphic-psychometric correlations within the larger SOL Sueño dataset (n=1761, Fig. S1), focusing on participant responses to the ESS, STAI-10 (Spielberger/State Trait Anxiety Inventory) and CESD-10 (Center for Epidemiological Studies Depression Scale). For each survey, we dichotomously assigned SOL subjects as “low” or “high” via median split (Fig. S1B). We then fit three independent logistic regression models using demographic (age, sex, BMI) and actigraphic (alpha, beta, acrophase, amplitude, F, average total daily “sleep”) predictor variables. Since shorter recording times can confound estimates of RAR robustness, we included recording duration as an additional predictor variable. As shown in Figure 4A, high STAI-10 (anxiety) scores in SOL subjects were significantly associated with female sex, a more delayed/left-shifted acrophase and lower F and amplitudes. When applied to PWE (Fig. 4B), the model displayed an accuracy of ∼70% for GAD-7 self-report, partitioning PWE into two clusters that also displayed contrasting scores on the QIDS-SR, PHQ-9 and AEP. Both groups were similar in BMI and were prescribed a similar landscape of major ASMs. The predicted-high anxiety group included significantly more female patients (χ^2^ = 5.5, p<0.05), and lower F scores in this subgroup were associated with *increased* intradaily variability (IV) and a trend for *decreased* interdaily stability (IS).

**Figure 4:**
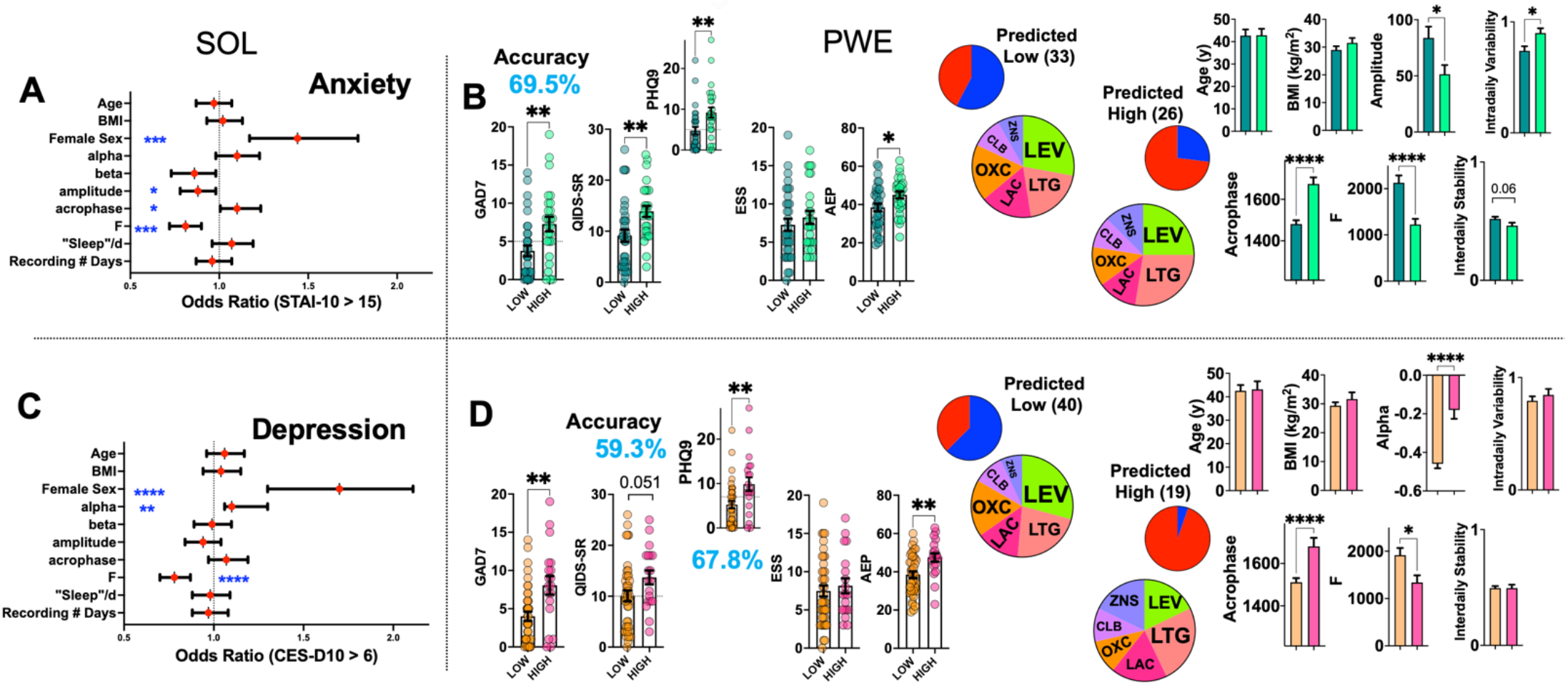
Models to Predict Psychiatric Comorbidity in PWE. A. Odds ratios and 95% confidence intervals (logistic regression) for predictor variables that determine greater than median STAI-10 scores (anxiety) within the SOL Sueño dataset. B. When applied as a classifier to PWE, this model classified subjects with an accuracy of 69.5%. C. Odds ratios and 95% confidence intervals for predictor variables that determine greater than median CESD-10 scores (depression) within the SOL Sueño dataset. D. When employed to classify PWE as low or high depression, the model displayed an accuracy of 59.3% (QIDS-SR scores) and 67.8% (PHQ-9 scores). Pie charts of ASM prescriptions reflect the 6 most commonly prescribed agents, including LEV (levetiracetam), LTG (lamotrigine), LAC (lacosamide), OXC (oxcarbazepine), ZNS (zonisamide) and CLB (clobazam). Mean ± s.e.m shown for all. *, **, **** indicating p < 0.05, p<0.01 and p<0.0001 respectively (unpaired two-tailed Student’s T test). ESS: Epworth Sleepiness Scale, AEP: Adverse Event Profile, PHQ: Patient Health questionnaire, GAD-7: Generalized anxiety disorder-7, QIDS-SR: Quick Inventory of Depression-Self Report.

High depression self-ratings in SOL subjects were also significantly associated with female sex and low F, as well as higher alpha (wider rest phase). Model accuracy in PWE subjects varied from ∼60% (QIDS-SR) to ∼70% (PHQ-9, Fig. 4D). PWE within the predicted-high depression group were almost exclusively female (χ^2^ = 17.21, p<0.001), displayed significantly greater GAD-7 and AEP scores, and were more likely to take lamotrigine and zonisamide, in place of levetiracetam and oxcarbazepine. Significantly lower average F scores were *not* associated with rhythm fragmentation, as measured by IV or IS. Both models outperformed an unsupervised classification model achieved via k-means clustering the same set of 10 demographic and actigraphic parameters. This approach (Fig. S2A) partitioned subgroups that displayed vastly different RAR patterns, but which were not significantly different in scores from all six psychometric instruments.

ESS was the sole assay identically employed in both cohorts, and scores were significantly higher in PWE as compared with matched SOL controls (Fig. S2B). High ESS scores in SOL Sueño subjects were significantly associated with low F and low alpha (wider active phase), but not female gender or total sleep duration. When applied to PWE, this model performed *worse* than chance (accuracy ∼40%, Fig. S2C). This suggests that unlike depression or anxiety symptoms, subjective sleepiness in PWE (i) may not be associated with definite RAR changes in PWE (Fig. 3D), or (ii) that RAR distortions related to sleepiness in PWE may be quite distinct from nonepileptic populations.

## Discussion

This report provides a cross-sectional evaluation of RARs in adult patients with focal epilepsy and examines whether variations in epilepsy severity and prevalent neuropsychiatric symptoms are associated with unique and stereotyped RAR variations. Our final sample is considerably smaller than many larger well-powered datasets (e.g., NHANES, SOL), but remains the largest outpatient actigraphic evaluation of PWE to date. Our cohort featured high rates of medically intractable epilepsy (2/3^rds^, compared with ∼1/3^rd^ observed in community estimates^68^), a subgroup prone to higher rates of psychiatric comorbidity and the behavioral side effects of ASMs^8,11,13,14,69^. RAR comparisons from within this cohort revealed that increases in seizure severity (and resultant ASM polytherapy) are associated with a dampened RAR amplitude and a wider rest phase, featuring more sleep. In contrast, prevalent anxiety and depression symptoms were associated with diminished RAR robustness and delayed acrophase, respectively. These initial results clearly deserve replication in larger multicenter cohorts. As it pertains to the benefits of wearable-based disease monitoring in PWE, our findings suggest that wrist accelerometry data may be combined with other continuously tallied physiological measures^70^ (recorded by the *same* device) to simultaneously ascertain convulsive seizure occurrence and appraise, or at least raise concern for, comorbid psychiatric symptoms.

Our conclusions come with the disclaimer, inherent to all cross-sectional actigraphic characterizations, that our detected associations cannot identify causal relationships. Thus (as an example), reductions in RAR amplitude in patients receiving ASM polytherapy may be the result of motor retardation imparted *by* ASMs, or an RAR trait that is biologically linked to drug refractoriness, or both. Our cross-sectional study design also limits the assessment of true biomarker potential, i.e., whether improvements in anxiety or depression over time are at all linked to a normalization of RAR structure. The inverse scenario also remains unanswered, i.e., whether behavior modifications designed to boost RAR robustness or alter acrophase can improve subjective measures of anxiety and depression^71^, and whether such improvements in psychiatric symptom burden may positively impact seizure frequency. Prospective clinical trials to answer these crucial questions could employ actigraphy not just as an objective endpoint, but also to assess adherence/compliance to treatment, akin to measuring serum ASM levels. Additional longitudinal studies may also assess whether specific RAR profiles can predict the likelihood of ASM side effects^72^.

The actigraphic-psychometric correlations that we compute from within the larger SOL Sueño dataset provide previously unrecognized insights into the connections between RAR robustness (F) and the dimensionalization of prevalent psychiatric symptoms. While low F scores in isolation may be diagnostically nonspecific, the presence and/or extent of any associated RAR changes may be able to potentially distinguish between symptom clusters. High *anxiety* scores were associated with a low F, delayed acrophase and reduced amplitude, while high *depression* scores were linked to a lower F and larger alpha values (wider rest phase). In contrast, low F and *low* alpha values significantly predicted high subjective sleepiness. Reductions in RAR robustness common to all three symptom clusters speak to the well-known relationships between mental health and the establishment of a daily routine^73,74^. One approach that automatically institutes a routine (at least on weekdays) is employment or fulltime education, often inaccessible to many PWE due to the burden of recurrent seizures, mild to moderate intellectual disability, driving restrictions or attention deficits. As expected, PWE subjects that were employed or enrolled in educational activities displayed a significantly *higher* F and amplitude, with lower QIDS-SR and AEP scores (Fig. S2D). This subgroup was also significantly younger, which may explain higher amplitudes but not higher F scores^39^.

As the ultimate test of our SOL-derived hypotheses, we asked whether a binary classifier derived from the richly annotated SOL dataset would have any success in predicting the burden of prevalent neuropsychiatric symptoms in our PWE population. There are many reasons why the accuracy of such a supervised classifier might have been poor. First, anxiety and depression surveys employed in both studies were distinct (STAI-10 vs GAD-7, CESD-10 vs QIDS-SR/PHQ-9), and to our knowledge, there are no controlled head-to-head comparisons of these instruments. Second, some have proposed that anxiety- and depression-related symptoms in PWE tend to occur in constellations that are unique to epilepsy (“*interictal dysphoric disorder*^75^”) and measured through distinct psychometric instruments, although this concept remains somewhat controversial^76^. Third, by definition, SOL subjects were self-identified Hispanic/Latino persons, while our PWE population was mixed in race/ethnicity, reflecting the cultural diversity of Houston, TX. Within the NHANES study^39^ at least, Hispanic subjects displayed a significantly higher amplitude and RAR robustness (compared with non-Hispanic whites, blacks and Asians), although it is not at all clear whether these differences reflect sociocultural constraints or a genetically mediated biological phenomenon. Fourth, the prevalence of epilepsy in the SOL population is unknown. Fifth, and perhaps most importantly, the sedation/hypoactivity/motor retardation imposed by ASMs may effectively obscure a classifier derived from nonepileptic populations and lead it to incorrectly recognize low RAR amplitude as a sign of depression and related syndromes. Despite these various caveats, we dichotomously classified anxiety and depression symptoms with an accuracy of approximately 70%. We take this optimistically as a sign that *epilepsy-specific* models derived from a larger, more diverse group of PWE (including patients with generalized epilepsy syndromes), may achieve even higher accuracy rates. The successful design of any such digital “phenotyper” will ideally have to overcome the central challenge of distinguishing between “psychomotor” retardation (as a sign of depressed mood) and motor retardation associated with ASM intake (“neuromotor?”). Indeed, as we continue to build a detailed understanding of disease-RAR relationships, it is already clear that specific RAR distortions are unlikely to provide any diagnostic specificity, and therefore must always be informed by clinical context. For example, the combination of lower F and lower amplitudes has been identified in studies of patients with depression^34-37^, obesity^40^ and late-stage Parkinson’s disease^77^.

For three PWE subjects in this study implanted with responsive neurostimulation (RNS) devices^64^, we were able to simultaneously juxtapose raw RARs with the timing of registered “long episodes” (sustained bursts of epileptiform activity) and neurostimulation treatments, both generally approximating the burden/severity of ongoing epileptiform activity. In two patients, clear circadian fluctuations in RNS treatments were evident (Fig. 5, A: diurnal, B: nocturnal). A third patient with nocturnal predominant RNS treatments (Fig. 5C) experienced a sustained increase in RNS treatments beginning on their third day of recording, together with an increase in measured long episodes. A closer review of the underlying actogram revealed that this change occurred following a night of prominent sleep deprivation/sleep delay. These examples suggest that at least in some patients, actigraphy may adjunct chronic ambulatory electrocorticography by straightforwardly depicting (in a patient-specific manner) how shifts in sleep patterns may dynamically impact seizure risk.

**Figure 5:**
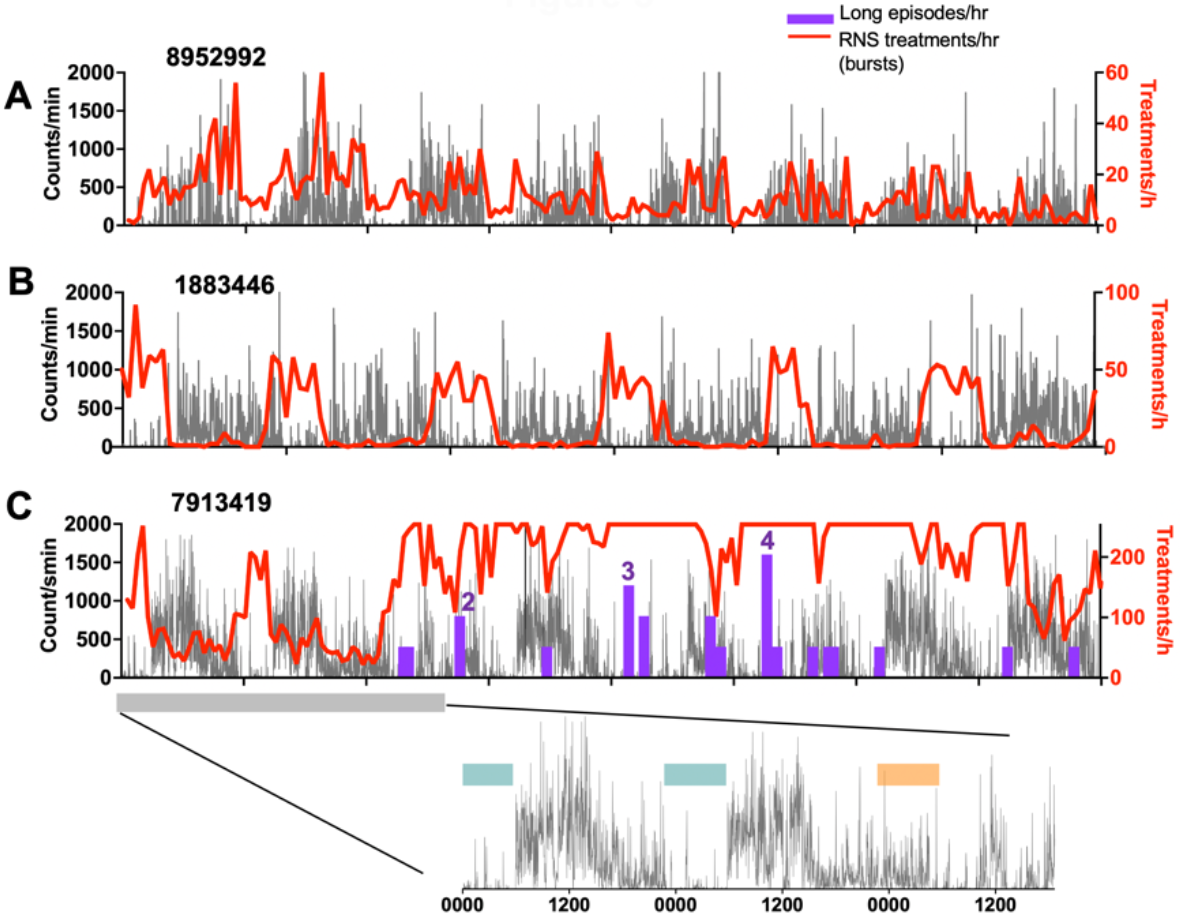
Actigraphy Combined with Responsive Neurostimulation (RNS). Superimposed actigraphy and RNS treatments illustrating examples of predominantly diurnal RNS treatments (A) and nocturnal treatments (B). In patient C, nocturnal predominant RNS treatments transitioned to a more sustained tonic rate of treatments beginning by Day 3. Inset: Raw actigraphy recordings confirmed that prior to this change, the patient experienced a night of poor sleep.

One of the most poorly recognized advantages of actigraphic endpoints lies in their ability to inform preclinical/basic science studies and enable more fluid bench-bedside translational efforts. With the increasing popularity of continuous, *in situ*, digital phenotyping tools in human subject research, prolonged continuous behavioral assessments of rodent or zebrafish models can now be seamlessly obtained through solutions that employ automated video tracking within the home-cage/-tank^29,78^. These technologies have been successfully employed to measure the severity of post-ictal locomotor suppression^79^, locomotor profiles associated with ASM exposure^80,81^ and model-specific distortions in rest/activity rhythms^82,83^. Importantly, such preclinical platforms can elaborate the locomotor consequences of ASM exposure in animals without *epilepsy*, while simultaneously assessing RARs in spontaneously seizing animal subjects in the *absence* of ASM exposure, thereby providing a fertile substrate to directly test causal hypotheses derived from human subject studies.

In conclusion, our study shows that actigraphy-derived determinations of RAR structure and timing in PWE may provide a range of objective endpoints to adjunct seizure diaries and survey-based assessments of wellbeing/quality of life. This approach may be particularly powerful in patients with disabling speech or language disturbances, and/or in those suspected to have comorbid insomnia, hypersomnia, or circadian rhythm sleep-wake disorder. Our results confirm the role of RAR robustness as a critical vital sign of mental wellbeing, and further, provide physiological “units of analysis”^84^ to uncover the neural circuits underlying anxiety and depression symptoms in transdiagnostic evaluations of patients with or without chronic medical comorbidities.

## Acknowledgements

VK acknowledges research funding from the NIH (1K08NS110924), the Mike Hogg Foundation and seed funding from the Baylor College of Medicine Office of Research.

---

**Supplemental Table 1.**
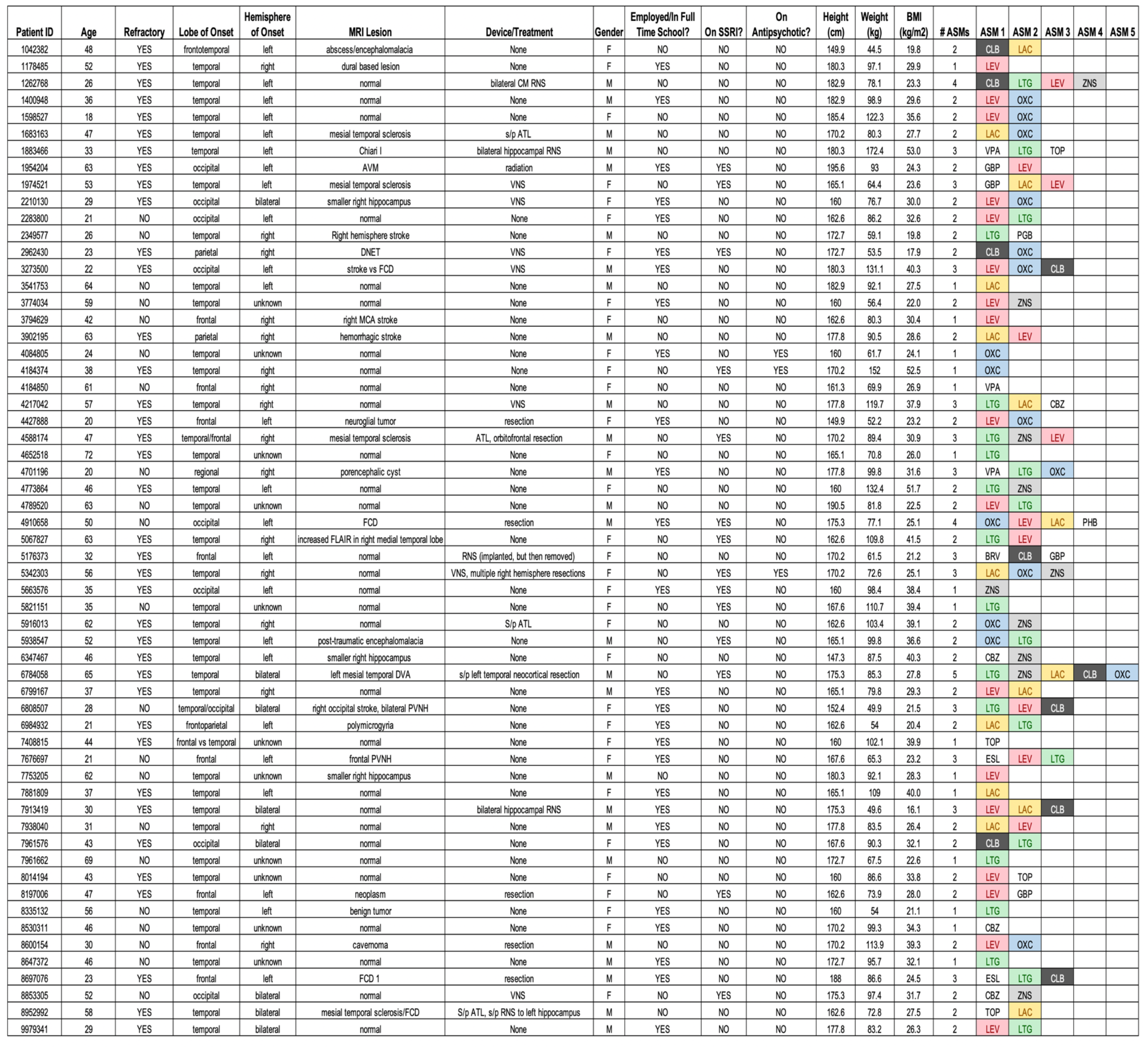
Abbreviations ATL: Anterior temporal lobectomy, ASM: Antiseizure medication, AVM: Arteriovenous malformation, DVA: Developmental venous anomaly, CM: centromedian thalamus, DNET: dysembryoplastic neuroepithelial tumor, RNS: Responsive neurostimulation (Neuropace), VNS: Vagus Nerve Stimulation, s/p: status post, PVNH: Periventricular nodular heterotopia, FCD: focal cortical dysplasia, CLB: Clobazam, LEV: Levetiracetam, LAC: Lacosamide, TOP: Topiramate, ZNS: Zonisamide, BRV: Brivaracetam, OXC: Oxcarbazepine, CBZ: Carbamazepine, ESL: Eslicarbazepine, CBZ: Carbamazepine, VPA: Valproic acid, GBP: Gabapentin, PGB: Pregabalin, LTG: Lamotrigine, PHB: Phenobarbital

**Supplemental Figure 1:**
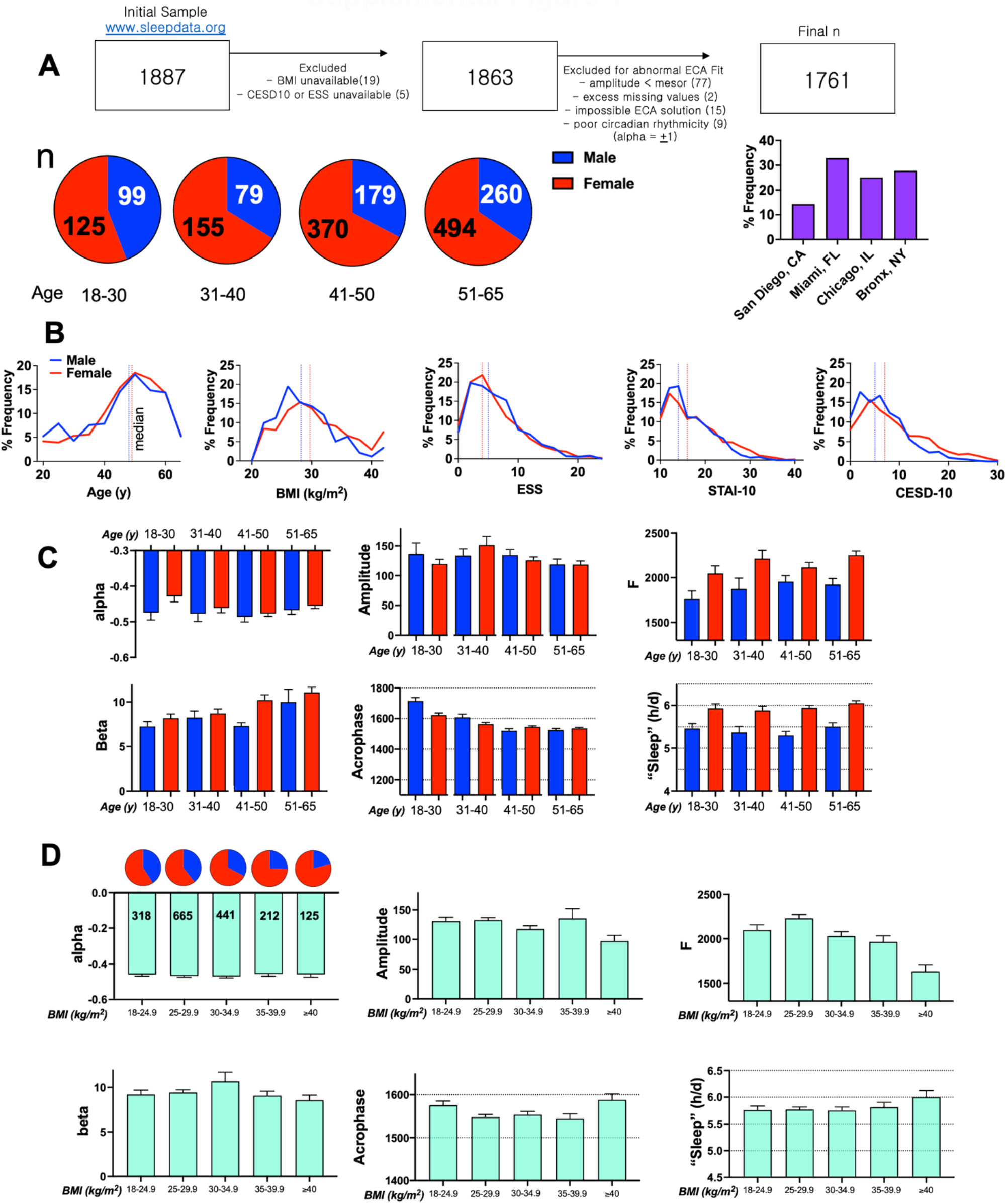
Demographic and RAR characteristics of the SOL Sueño actigraphy dataset. A: Summary of excluded subjects. Below, sex breakdown by age-group, and right: city of recruitment. B. Frequency distribution histograms with delineated median (blue – male, red – female). C. Across age groups, female subjects displayed higher alpha (*sex*, F_1,1753_ = 4.5, p<0.05), more sleep (*sex*, F_1,1753_ = 57.1, p< 0.0001 and higher F (*sex*, F_11,1753_ = 22.39, p<0.001). Beta increased with age but not sex (*age*, F_3, 1753_ = 3.79, p<0.05). Acrophase varied by both sex and age (*sex x age*, F_3,1753_ = 21.96, p<0.0001), and amplitude did not show any age- or sex-related variations (p>0.05). D. RAR amplitude decreased with increasing BMI (F_4,1756_ = 2.6, p<0.05), while acrophase changed in a somewhat U-shape manner (F_4,1756_ = 3.19, p<0.05). Higher BMIs are associated with substantially lower F values (F_4,1756_ = 10.2, p<0.001), while significant BMI-dependent changes in sleep were not observed. Mean ± s.e.m shown for C and D.

**Supplemental Figure 2:**
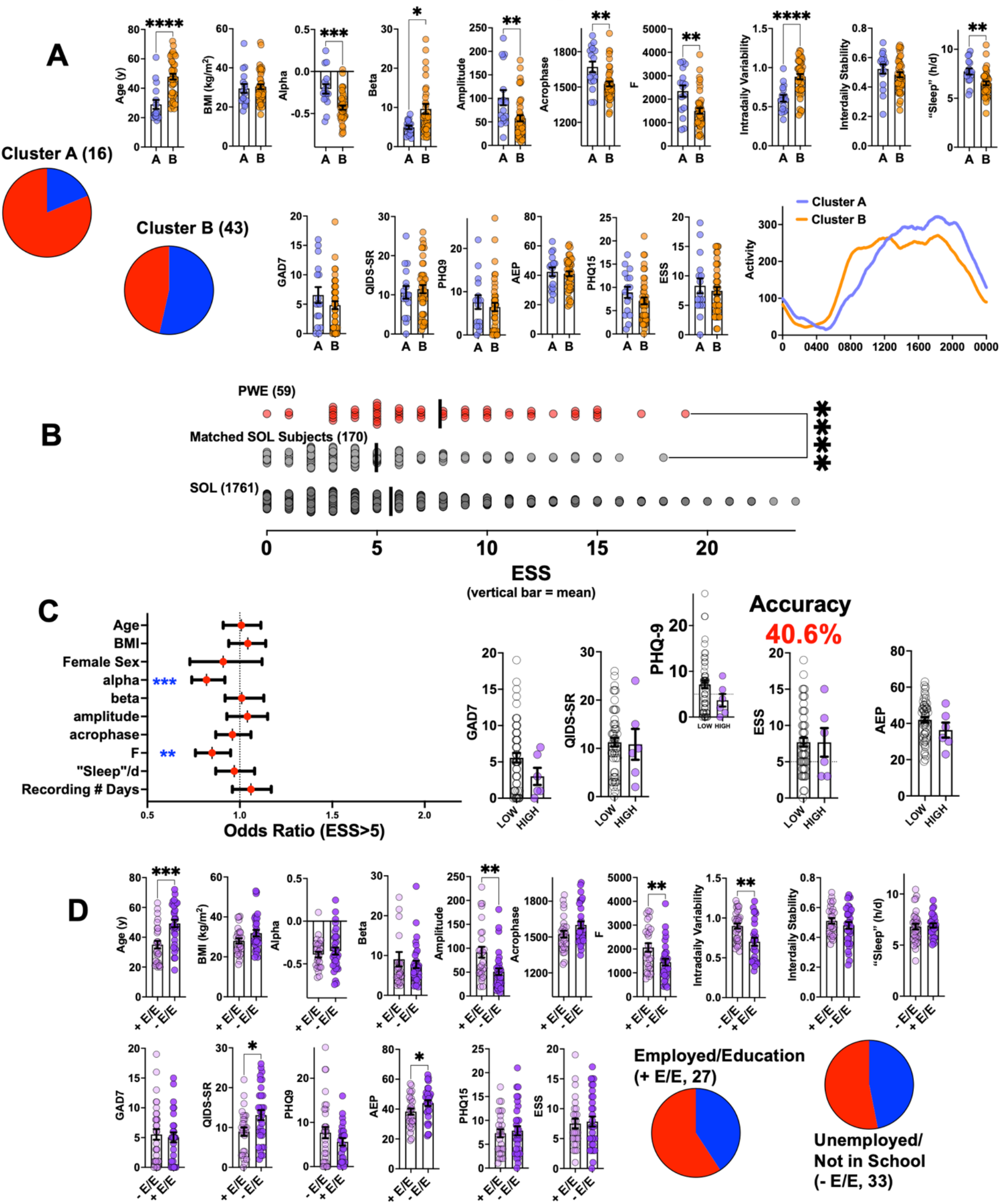
A: Partitioning PWE into two subgroups via K-means clustering (based solely on demographic and actigraphic parameters) distinguished a smaller, younger and mostly female cluster (A) from an older and larger cluster (B) with a similar number of male and female subjects. Patients in cluster B displayed a lower amplitude, alpha, F and lower amounts of sleep, together with a higher beta (increased steepness). However, clusters A and B were largely similar across psychometric indices. B. Distribution of total ESS responses across PWE, SOL matched subjects (Fig. 1) and the larger SOL dataset. C. Odds ratios and 95% confidence intervals (logistic regression) for predictor variables that determine greater than median ESS scores (sleepiness) within the SOL Sueño dataset. B. When applied as a classifier to PWE, this model performed poorly, classifying subjects with an accuracy < 50%. D. PWE subjects that were either employed or in educational activities were on average significantly younger, displayed higher amplitudes and F, in concert with lower QIDS-SR and AEP scores. Mean ± s.e.m shown for all. *, **, ***, **** indicating p < 0.05, p<0.01, p<0.001 and p<0.0001 respectively (unpaired two-tailed Student’s T test). ESS: Epworth Sleepiness Scale, AEP: Adverse Event Profile, PHQ: Patient Health questionnaire, GAD-7: Generalized anxiety disorder-7, QIDS-SR: Quick Inventory of Depression-Self Report.

